# Land use effects on soil microbiome composition and traits with consequences for its ecosystem carbon use efficiency

**DOI:** 10.1101/2024.04.05.588235

**Authors:** Lisa Cole, Tim Goodall, Nico Jehmlich, Robert I. Griffiths, Gerd Gleixner, Cecile Gubry-Rangin, Ashish A. Malik

## Abstract

The soil microbiome determines the fate of belowground inputs of plant fixed carbon. The shifts in soil properties caused by changes in land use leads to modifications in microbiome structure and function, resulting in either loss or gain of soil organic carbon (SOC). Soil pH is the primary factor regulating microbiome characteristics leading to distinct pathways of microbial carbon cycling, but the underlying mechanisms remain understudied. Here, the taxa-trait relationships behind the variable fate of SOC were investigated across two temperate paired land use intensity contrasts with differing soil pH using metaproteomics, metabarcoding and a ^13^C labelled litter decomposition experiment. ^13^C incorporation into microbial biomass increased with land use intensification in low pH soils but decreased in high pH soils, impacting ecosystem carbon use efficiency (CUE) in opposing directions. Reduction in biosynthesis traits across land use intensity contrasts was due to increased abundance of proteins linked to resource acquisition and stress tolerance. These community-level trait trade-offs were underpinned by land use intensification-induced changes in dominant taxa with distinct traits. These trait changes alter the balance of decomposition and stabilisation of carbon in soil through divergent pH-controlled pathways. In low pH soils, land use intensification alleviates microbial abiotic stress resulting in increased CUE but promotes decomposition and SOC loss. In contrast, in high pH soils, land use intensification increases microbial physiological constraints and decreases CUE, leading to reduced necromass build-up and SOC stabilisation. We demonstrate how microbial CUE can be decoupled from SOC highlighting the need for its careful consideration in predicting or managing SOC storage for soil health and climate change mitigation.

## Introduction

Soils are under pressure to deliver multiple ecosystem services, especially food production. This has led to the expansion of agriculture into pristine environments and increased intensification. There is a growing recognition that the intensive use of soils is detrimental to soil health, changing soils’ inherent biodiversity and risking the services that they provide [1, 2]. The world’s soils have historically lost 133 Pg of carbon due to land use intensification [3]. However, degraded soils low in organic matter also represent an opportunity to adopt regenerative management promoting soil carbon storage that may help mitigate this issue [1, 4, 5]. To better achieve this aim, it is vital to understand the role of soil microbes in carbon cycling, as the microbiome is the gatekeeper of soil-atmosphere carbon exchange controlling the fate of carbon in soils [6].

A new paradigm recognises the direct, significant contribution of microbes in transforming photosynthetically derived carbon into soil organic carbon [7], by stabilising dead microbial biomass (necromass) onto mineral surfaces to enable persistent, long-term carbon storage [8]. Microbial CUE is a vital ecosystem trait that determines soils’ ability to accumulate carbon [9] and is measured as the incorporation of organic carbon from the environment into its biomass through growth [10, 11]. A higher microbial CUE implies more efficient biomass production and a lower respiratory loss [12]. Increased growth and death of microbes results in a bigger necromass pool that on association to mineral surfaces can form persistent SOC thereby promoting soil carbon storage [13]. As microbes become more efficient in using carbon, higher carbon storage is observed in soils, a pattern that is detectable at the global scale [14]. Increased microbial CUE therefore offers the potential to increase the necromass pool for stabilisation in the mineral-associated organic matter resulting in long term SOC storage.

Microbiome diversity and function are responsive to environmental gradients [15–17] and microbial biomass is generally greatest under low intensity land use [18, 19]. Increased necromass production for promoting soil carbon storage is likely best achieved at grassland sites with low land use intensification. Given the degraded state of many of the world’s agricultural soils that have lost SOC, croplands represent a habitat where carbon storage could be promoted through microbiome-mediated processes. Therefore, it is crucial to understand how land use intensification impacts key microbial traits such as CUE [20–22]. This knowledge would enable us to better manage degraded soils to enhance microbial CUE and promote SOC stabilisation, providing many benefits for soil health, soil biodiversity, and climate change mitigation [4, 23, 24].

A positive relationship between microbial biomass and SOC concentration has been observed across 21 paired land use contrasts in the UK [25]. However, land use intensification effects on community-level CUE were complex and were better explained through interactions of multiple soil properties. Of these, soil pH was identified as the dominant factor, as converting grasslands to cropland tends to increase soil pH [25]. Soil pH has been previously found to be the main factor influencing soil microbial diversity [26, 27]. The UK-wide study suggested two distinct, pH-dependent mechanisms of soil carbon accumulation [25]. Acid, wet, and anoxic conditions limit microbial growth and decomposition [25], accumulating part-decomposed plant materials at the soil surface and high SOC in the upper horizons [28]. In contrast, well-drained neutral to alkaline pH soils provide conditions more conducive to microbial growth, promoting necromass generation for stabilisation as SOC [25]. Thus, soil pH can be used as a proxy to study the divergent effect of land use intensification on soil microbiomes and carbon cycling.

The trait-based life history strategies of the resident microbiome can explain the divergent mechanisms of microbial SOC accumulation. A life history framework has been proposed for microbes classifying them into three main strategies: high yield (Y), resource acquisition (A) and stress tolerance (S) with multiple underlying traits [29]. These traits correlate due to physiological or evolutionary trade-offs, influenced by the environmental conditions such as resource availability and abiotic stress [25, 29]. In low resource environments, typical of high land use intensity soils (e.g. arable systems where plant biomass inputs to soil tend to be low), traits that enable microbial survival and activity include investment into the production of extracellular enzymes for resource acquisition pathways [29, 30]. In soils under high land use intensification, microbes are exposed to increased frequency of drought stress as tillage leads to soil aggregate disruption and lower water holding capacity [31]. Investment in stress tolerance in high land use intensity soils can often be observed with chaperone proteins such as Chaperonin GroEl that prevent stress-induced misfolding of proteins [25]. These increased cellular investments into stress alleviation and resource acquisition trade off with microbial growth yield due to the diversion of resources from growth and biosynthesis. The reduced biomass (and subsequent necromass pool) and the increased respiratory loss reflect lower SOC accumulation rates [25, 29]. Furthermore, under intense abiotic pressure, such as drought, microbes might also shift to a dormancy state, reducing ecosystem CUE [32, 33].

While microbial community-level traits such as CUE have been linked to ecosystem measures such as changes in SOC, identifying taxonomic groups contributing to higher CUE is challenging. Previous studies have aimed to do this, by assigning microbial taxa to trophic groups or life history strategies such as the copiotroph-oligotroph dichotomy [34, 35]. It was observed that copiotrophs invest in a competitive strategy and have a high maintenance respiration, which reduces their CUE. In contrast, oligotrophs maintain growth over respiration in low quality resource environments, thereby increasing their CUE [35, 36]. However, the copiotroph-oligotroph dichotomy does not exist at broader levels of taxonomic linages [37]. Therefore, linking a comprehensive set of traits (such as those for Y-A-S life history strategies) to taxonomic identity is essential to better understand how organismal physiology influences ecosystem-level processes.

This study investigated the microbial community response to land use intensification in two temperate sites of contrasting soil pH to understand how taxonomic and trait shifts impact soil carbon cycling. Our current understanding of the microbial traits underpinning SOC stabilisation processes is mainly obtained through analysing a community response, often using an emergent trait such as CUE. In addition to this approach, we aim to identify how changes in the abundance of dominant microbial taxa caused by land use intensification led to shifts in key microbial traits, emergent ecosystem CUE, and SOC decomposition and stabilisation rates. We hypothesise that increased land use intensification impacts soil properties, with a shift from high growth yield taxa to resource acquiring and stress tolerant taxa in the microbial community, resulting in lower CUE and SOC stabilisation. Using metaproteomics and metabarcoding, we identified the dominant taxonomic groups with different Y-A-S traits and related them to ecosystem CUE measures. Therefore, this study demonstrates how land use intensification selects microbial communities with variable organismal traits impacting the soil carbon cycling.

## Methods

### Site description

To understand how microbial taxonomy and traits influence soil carbon dynamics in soils of differing land use intensity, we chose two sites with contrasting pH that were previously studied as part of a landscape scale survey [25]. The low pH site (pH 5.2), Kirkton located in Perthshire (Table 1), has historically undisturbed plots representative of wet acid upland podzols [28] with high SOC in the upper horizons. These soils are dominated by U4d (*Festuca ovina–Agrostis capillaris*–*Galium saxatile, Luzula multiflora–Rhytidiadelphus loreus* subcommunity) and the U5a (*Nardus stricta–Galium saxatile* species-poor subcommunity) grasslands of the National Vegetation Classification [38]. The contrasting plot has soils improved to support agricultural activities by drainage and liming, this raised soil pH to 6.4. The high pH site (pH 7.7) was a chalk grassland at Parsonage Down National Nature Reserve located in Wiltshire (Table 1). The low land use intensity plot has not been ploughed in the last 100 years and supports a herb-rich plant community dominated by CG2 *Festuca ovina–Avenula pratensis* grassland [38]. The high land use intensity arable cropland plot at this site has a soil pH of pH 8 (Table 1). In both sites, land use intensification led to a loss of soil organic carbon. Pairwise t-test was performed to ascertain the effect on land use intensity on soil properties.

**Table 1:** Site characteristics given as mean values (± standard error) with statistical comparison of land use intensity contrasts within sites given by p values of t-test.

|  | Low pH site: Perthshire, UK |  |  | High pH site: Wiltshire, UK |  |  |
| --- | --- | --- | --- | --- | --- | --- |
| Land use intensity | Low | High | Pairwise t-test p-value | Low | High | Pairwise t-test p-value |
| Land management | Unimproved grassland | Intensive grassland |  | Unimproved grassland | Intensive arable |  |
| Soil pH | 5.2 ($\pm 0.2$) | 6.4 ($\pm 0.1$) | 0.011 | 7.7 ($\pm 0.03$) | 8.0 ($\pm 0.04$) | 0.001 |
| Soil C (%) | 23.8($\pm 8.5$) | 4.3 ($\pm 0.6$) | 0.145 | 10.4 ($\pm 0.5$) | 3.8 ($\pm 0.1$) | 0.004 |
| Moisture (%) | 72.1 ($\pm 13.6$) | 41.7 ($\pm 2.5$) | 0.153 | 43.1 ($\pm 4.9$) | 30.4 ($\pm 4.9$) | <0.001 |

### Experimental design

At each plot, three spatially dispersed soil cores (5 cm diameter, 15 cm deep) were sampled. Soil samples were preserved at 4°C following removal of vegetation and homogenisation by sieving (< 4mm). Mesocosms were established in Petri dish plates containing 10 g (dry weight equivalent) soil, maintained at field moisture gravimetrically and incubated at 21°C for 7 days. After this time, 3 mg ^13^C-labeled *Chenopodium* sp. leaf litter was mixed thoroughly with the soil in each mesocosm (n=3). As the amount of carbon in the added litter was very low (<1%) relative to the existing soil carbon, the influence of litter addition on microbial community taxonomy and function is considered negligible. The ^13^C-labeled leaf litter was produced by growing *Chenopodium* sp. in a closed chamber containing ∼1 atom% ^13^C-CO_2_ at a concentration of 400 ppm, followed by drying of leaves and homogenisation by grinding. Mesocosms were destructively harvested on day 0 (just before litter addition) and days 2, 8 and 36 following litter addition. ^13^C-labelling of the litter enabled ^13^C to be traced into separate pools as microbial biomass, respired CO_2_, and bulk soil. The labelled substrate was added at a single time, allowing the monitoring of the microbial ecosystem CUE over the incubation period.

### Microbial CUE

An aliquot (1 g) of the soil collected at each sampling point was placed in a sealed 10-ml glass vial with rubber septa and incubated overnight (for ∼16 h) at 21 °C in the dark to collect respired CO_2_ in the headspace. Concentrations of CO_2_ and its ^13^C content was analysed by gas chromatography isotope ratio mass spectrometer (GC-IRMS, Delta + XL, Thermo Fisher Scientific, Germany) coupled to a PAL autosampler (CTC Analytics) with general purpose (GP) interface (Thermo Fisher Scientific, Germany). DNA was extracted from 0.25 g soil at each sampling point using the PowerSoil-htp 96-well soil DNA isolation kit per manufacturer’s instructions (MO BIO Laboratories, UK) and its quality was checked by Nanodrop. Total extractable DNA concentration was also measured using a Qubit fluorometer, providing a proxy for microbial biomass. ^13^C content of DNA extracts was analysed by liquid chromatography isotope ratio mass spectrometer LC-IRMS (HPLC system coupled to a Delta + XP IRMS through an LC IsoLink interface; Thermo Fisher Scientific, Germany). This approach enabled quantification of the proportion of ^13^C labelled plant litter in total microbial DNA and respired CO_2_ during the incubation. Microbial CUE was calculated using the following equation:

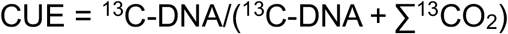

where ∑^13^CO_2_ is the cumulative litter-derived ^13^C lost during respiration. Statistical analyses and visualisations in ggplot2 [39] were performed using R software 2023.3.0 [40]. Multi-factorial ANOVA was performed to ascertain the effect of site, land use intensity and sampling time on ^13^C in DNA, ^13^C in respired CO_2_ and ecosystem CUE.

### Metabarcoding

DNA was extracted as described above. Amplicon libraries were constructed according to a dual indexing strategy [41] with each primer consisting of the appropriate Illumina adapter, 8-nt index sequence, a 10-nt pad sequence, a 2-nt linker and the amplicon specific primer. For prokaryotes, the V3-V4 16S rRNA amplicon primers from Kozich et al. [41] were used (CCTACGGGAGGCAGCAG and GCTATTGGAGCTGGAATTAC), for eukaryotes the 18S rRNA amplicon primers from Baldwin et al. [42] were used (AACCTGGTTGATCCTGCCAGT and GCTATTGGAGCTGGAATTAC). Amplicons were generated using a high-fidelity DNA polymerase (Q5 Taq, New England Biolabs). After an initial denaturation at 95 ºC for 2 minutes PCR conditions were: denaturation at 95 ºC for 15 seconds; annealing at temperatures 55 ºC, 57 ºC for 16S, 18S reactions respectively; annealing times were 30 seconds with extension at 72 ºC for 30 seconds; cycle numbers were 25 for 16S, and 30 for 18S; final extension of 10 minutes at 72 ºC was included. Amplicon sizes were determined using an Agilent 2200 TapeStation system and libraries normalized using SequalPrep Normalization Plate Kit (Thermo Fisher Scientific) and quantified using Qubit dsDNA HS kit (Thermo Fisher Scientific). Each amplicon library was sequenced separately on Illumina MiSeq using V3 600 cycle reagents at concentrations of 8 pM with a 5% PhiX Illumina control library. Raw data have been deposited in NCBI SRA under accession PRJNA1088078.

Sequenced paired-end reads were joined using PEAR[43], quality filtered using FASTX tools (hannonlab.cshl.edu), length filtered with the minimum length of 300 bp, presence of PhiX and adaptors were checked and removed with BBTools (jgi.doe.gov/data-and-tools/bbtools/), and chimeras were identified and removed with VSEARCH [44] using Greengenes 13_5 [45] and SILVA 132 [46] databases for 16S and 18S respectively (at 97%). Singletons were removed and the resulting sequences were clustered into operational taxonomic units (OTUs) with VSEARCH at 97% sequence identity. Representative sequences for each OTU were taxonomically assigned by RDP Classifier [47] with the bootstrap threshold of 0.8 or greater using the Greengenes 13_5 and SILVA 132 databases (16S and 18S respectively) as the reference. Unless stated otherwise, default parameters were used for the steps listed. Taxonomic groupings of prokaryotes were presented using the older taxonomic classification to compare with proteomics-derived taxonomy. Only three major groups of eukaryotes: fungi, Ciliophora and Cercozoa were analysed. α-diversity indices (Shannon Weiner diversity index, Pielou’s Evenness Index and OTU Richness) were calculated on rarefied data (2000 reads) using the vegan package in R [48] and visualisations were performed using ggplot2 [39]. β-diversity was assessed by in non-metric multidimensional scaling ordinations and running Permutational Multivariate Analysis of Variance (PERMANOVA) using vegan’s adonis2 function. Multi-factorial ANOVA was performed to ascertain the effect of site and land use intensity on diversity indices and the abundance of taxonomic groups of interest.

### Metaproteomics

Metaproteomic analysis was performed on soil microbial communities for day 0 and day 8 samples. Proteins were extracted from 5 g of soil (with two technical replicates) using the SDS buffer–phenol extraction method, followed by purification with 1D SDS-PAGE. The resultant product was subjected to tryptic digestion. Proteolytically cleaved peptides were separated prior to mass spectrometric analyses by reverse-phase nano HPLC on a nano-HPLC system (Ultimate 3000 RSLC nano system, Thermo Fisher Scientific, San Jose, CA, USA) coupled online for analysis with a Q Exactive HF mass spectrometer (Thermo Fisher Scientific, San Jose, CA, USA) equipped with a nano electrospray ion source (Advion Triversa Nanomate, Ithaca, NY, USA). Raw data were searched using Proteome Discoverer v1.4.1.14 (Thermo Fisher Scientific) against a FASTA-formatted database (Uniprot 05/2016) using the SEQUEST HT algorithm. Additional details on quality control, database searches, and filtering are described elsewhere [25]. The mass spectrometry data are available in the ProteomeXchange Consortium via the PRIDE partner repository with the identifier PXD010526. Functional annotation was performed using KEGG classifier and GhostKoala. Taxonomic origin was assigned to proteins using Unipept v3.2, enabling us to make function-taxonomy linkages. Two-factorial ANOVA was performed to ascertain the effect of site and land use intensity on proteomics-derived functional diversity index. Pairwise Indicator Species Analysis was performed to identify the protein functions that were significantly enriched in low- and high-intensity land use treatments at each site [25]. The abundance of different protein functions that were identified was then investigated in each taxonomic group of interest and this was plotted using ggplot2 by combining the geom_tile and geom_point functions. Pairwise t-test was performed to ascertain the effect on land use intensity on the abundance of protein functions under each taxonomic group.

## Results and discussion

### Land use intensification alters soil physicochemical properties

Land use intensification had profound effects on soil properties, significantly increasing soil pH at both sites (Table 1). At the low pH site, pH increased from 5.2 to 6.4 through liming that is necessary to achieve the optimum soil pH range for crop plant nutrient availability [49]. Improved drainage and crop cultivation reduced the soil moisture into an optimal range (40-50%), hence reducing anoxia, further alleviating physiological constraints on the soil microbiome. Thus, the wider assumption that land use intensification causes aridity and drought stress in soil microbiomes does not apply to poorly drained acidic soils [25]. Land use intensification only marginally increased soil pH at the high pH site – a shift of 0.3 units. These soils are inherently alkaline, and do not require pH adjustment through liming to support agriculture. Increased land use intensification at this site reduced soil moisture below the optimum range (40-50%), possibly increasing the risk of drought stress [50]. Therefore, soil conditions under increased land use intensification are likely more challenging for the microbiome at the tested high pH site.

Land use intensification at the tested low pH site resulted in over 80% of the SOC being lost relative to the unimproved soil (Table 1). Increased decomposition in organic soils under land use intensification is a key mechanism for SOC loss, as the carbon at these sites is particularly vulnerable to loss due to a lower proportion of mineral-associated organic matter or MAOM [4, 33, 51]. Land use intensification at our high pH site led to a marked SOC decline from 10.4% to 3.8% (Table 1), confirming that cultivated soils are prone to SOC loss [52]. These impacts of land use intensification on soil properties are in accordance with Malik et al. [25] who also observed increased soil pH under intensification that led to SOC loss and reduced soil moisture availability, being most pronounced in low pH soils.

### Land use intensification influences microbial growth, respiration, and ecosystem CUE

Microbial growth depended on land use intensification and site (Fig. 1a), with increased land use intensification resulting in an increase of ^13^C in the microbial DNA at the low pH site (54% more in the high than in the low land use intensity soil) and in a reduction of ^13^C in microbial DNA at the high pH site (35% less in the high than in the low land use intensity soil). This contrasting effect of land use intensity at the two sites is highlighted by the significant interactive effect of site and land use intensity (ANOVA, p<0.001). This result supports our hypothesis that land use intensification reduces carbon incorporation into microbial biomass, but only at the high pH site where land use intensification reduced soil resource availability and moisture. In contrast, land use intensification at low pH alleviated physiological constraints of acidity, wetness, and anoxia, enabling increased growth (Fig.1a).

**Fig. 1:**
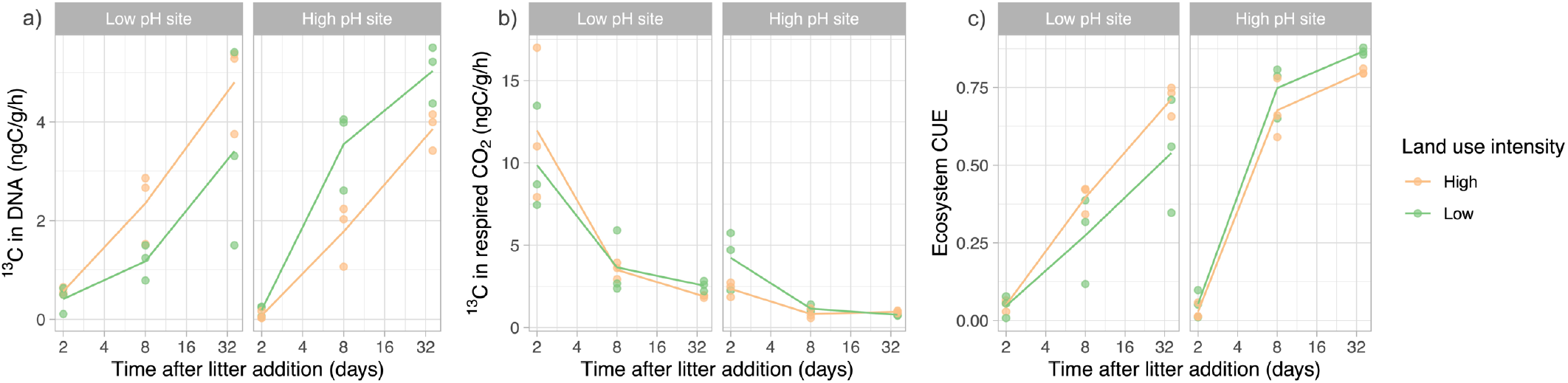
Isotopic incorporation from labelled litter into DNA as a proxy for microbial biomass (a) and respired CO_2_ (b). Patterns of estimated ecosystem CUE (c) under low (green) and high (orange) land use intensity. Points indicate individual samples, and the lines connect the mean values at each sampling time. ANOVA p values of the influencing factors of site (S), land use intensity (L) and their interaction (S×L) for (a) ^13^C in DNA - S: 0.37, L: 0.92, S×L: 0.006; (b) ^13^C in respired CO_2_ - S: <0.001, L: 0.95, S×L: 0.64; (c) CUE - S: 0.014, L: 0.91, S×L: 0.15.

We hypothesised that land use intensification results in an increase in the decomposition rate of an added complex resource, measured by an increased ^13^CO_2_ production. However, there was no difference in respiratory rate of the ^13^C labelled substrates in soils across the land use intensity contrasts at both sites (Fig. 1b).

Ecosystem-level microbial CUE deduced from ^13^C in the microbial DNA and respired CO_2_ was not statistically significant across land use intensity treatments, but the patterns suggest a reduction with land use intensification at the high pH site and the opposite at the low pH site (Fig. 1c). A similar pattern of community-level CUE has been previously observed in the same soils [25]. The increased CUE values over time following labelled litter addition highlights the long-term persistence of carbon in the microbial biomass due to substrate recycling in the microbial food web. Such measurements are key to studying the longer-term effects of microbial processes on soil carbon cycling; CUE measured over a longer incubation period (several weeks) enables inferring the complex interactions within the microbial community and between the microbial community and its abiotic environment [9]. The reduction in ecosystem CUE with land use intensification at the high pH site translates into lower biomass and necromass production with a lower SOC stabilisation potential [13, 53]. Conversely, land use intensification alleviated environmental stressors on the soil microbiome in low pH soil, promoting microbial growth. Here SOC change is decoupled from microbial CUE, and other biogeochemical mechanisms might be more important in controlling the rate of SOC loss or accumulation. It also highlights that current microbial CUE measurements do not always link to historical soil carbon changes. Therefore, future research must consider the balance between the biogeochemical processes of decomposition and stabilisation, including abiotic factors such as organic matter access, chemistry, and mineral stabilisation, when studying the impact of long-term land use change on changes in soil carbon storage.

### Land use intensification changes microbial diversity

The functional (inferred from metaproteomics) and taxonomic (inferred from metabarcoding) composition of microbial communities changed with land use intensification in the two sites (Fig. 2a-c). The sites strongly differed in microbial functional and taxonomic diversity (Fig. 2a-c), but the functional alpha diversity was not different across the land use intensity treatments at both sites (Fig. 2d). The community shifts over time were insignificant, suggesting that the small amount of plant litter that was added caused only minor changes in microbial taxonomy and function; all sampling points were therefore considered replicates to study the effect of land use intensification.

**Fig. 2:**
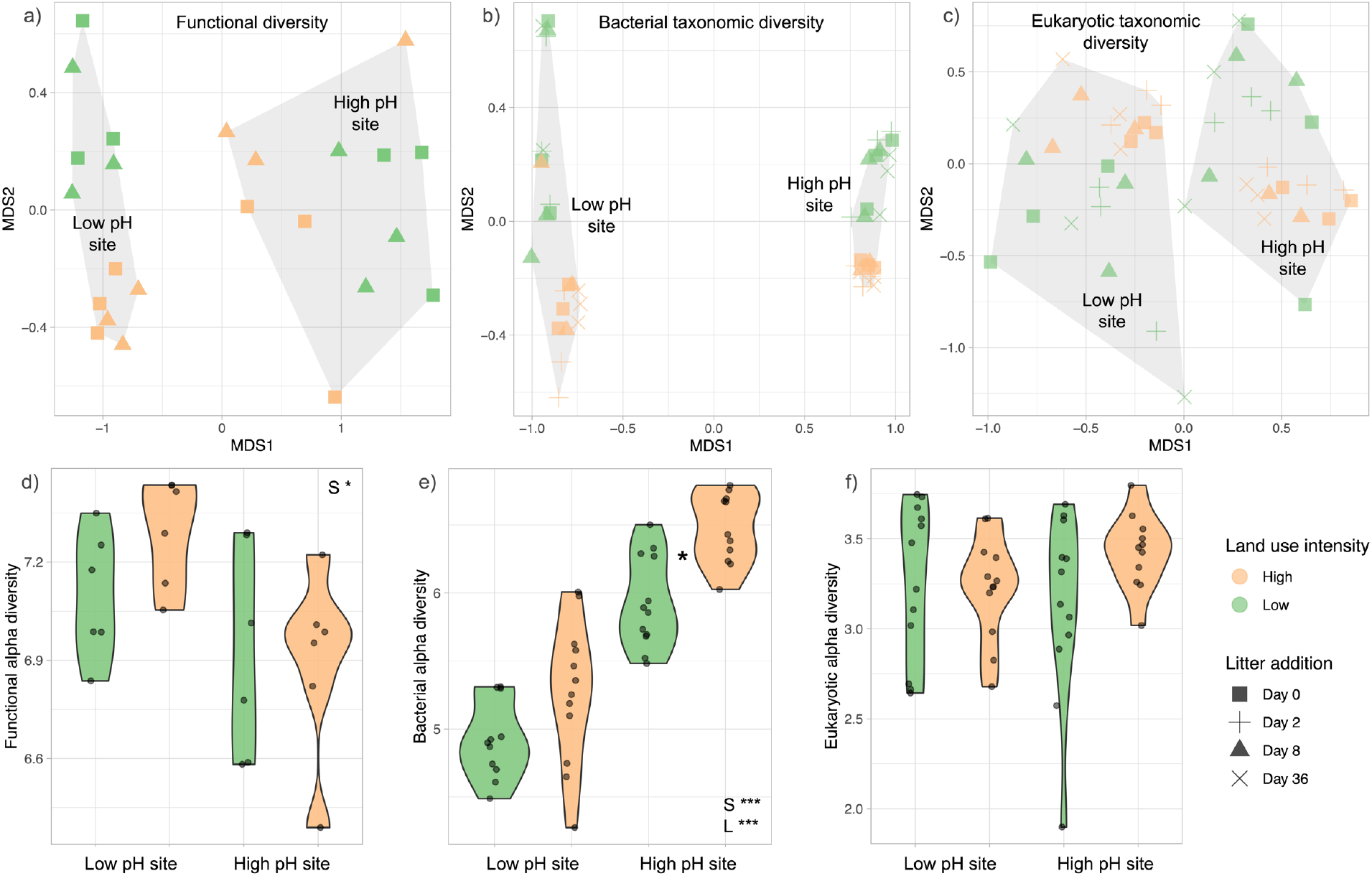
Ordination using nonmetric multidimensional scaling (NMDS) of metaproteomics-derived functions (a), 16S rRNA gene-derived bacterial taxonomy (b) and 18S rRNA gene-derived eukaryotic taxonomy (c). Similarly, Shannon’s diversity index was used to visualise functional alpha diversity (d), bacterial alpha diversity (e) and eukaryotic alpha diversity (f) under high (orange) and low (green) land use intensity at the two sites under study. In d-f, the presence of an asterisk between low and high land use intensity violins suggests statistically significant pairwise differences. Also displayed within d-f are statistically significant ANOVA results of the influencing factors of site (S), land use intensity (L) and their interaction (S×L); *** p < 0.001, ** p < 0.01, * p < 0.05 (non-significant results are not displayed). Note that metaproteomics was performed only at day 0 and day 8 after litter addition.

Bacterial alpha diversity significantly increased with land use intensification (Fig. 2e) which corroborates previously observed high bacterial diversity in agricultural soils [26, 54], contradicting the notion that disturbance decreases biodiversity [51]. Several explanations for this apparent paradox have been proposed, such as agricultural rotations increasing resource heterogeneity [55] and tillage redistributing plant litter to depth facilitating access to resources and growth to a diverse range of bacteria [56]. The high diversity of microbial taxa in agricultural soils could also represent relic DNA from dead microbes that sticks to soil minerals [57]. In contrast, eukaryotic alpha diversity (Fig. 2f) and the abundance of OTUs representing fungal taxa (Fig. 3j) were unaffected by land use intensification.

**Fig. 3:**
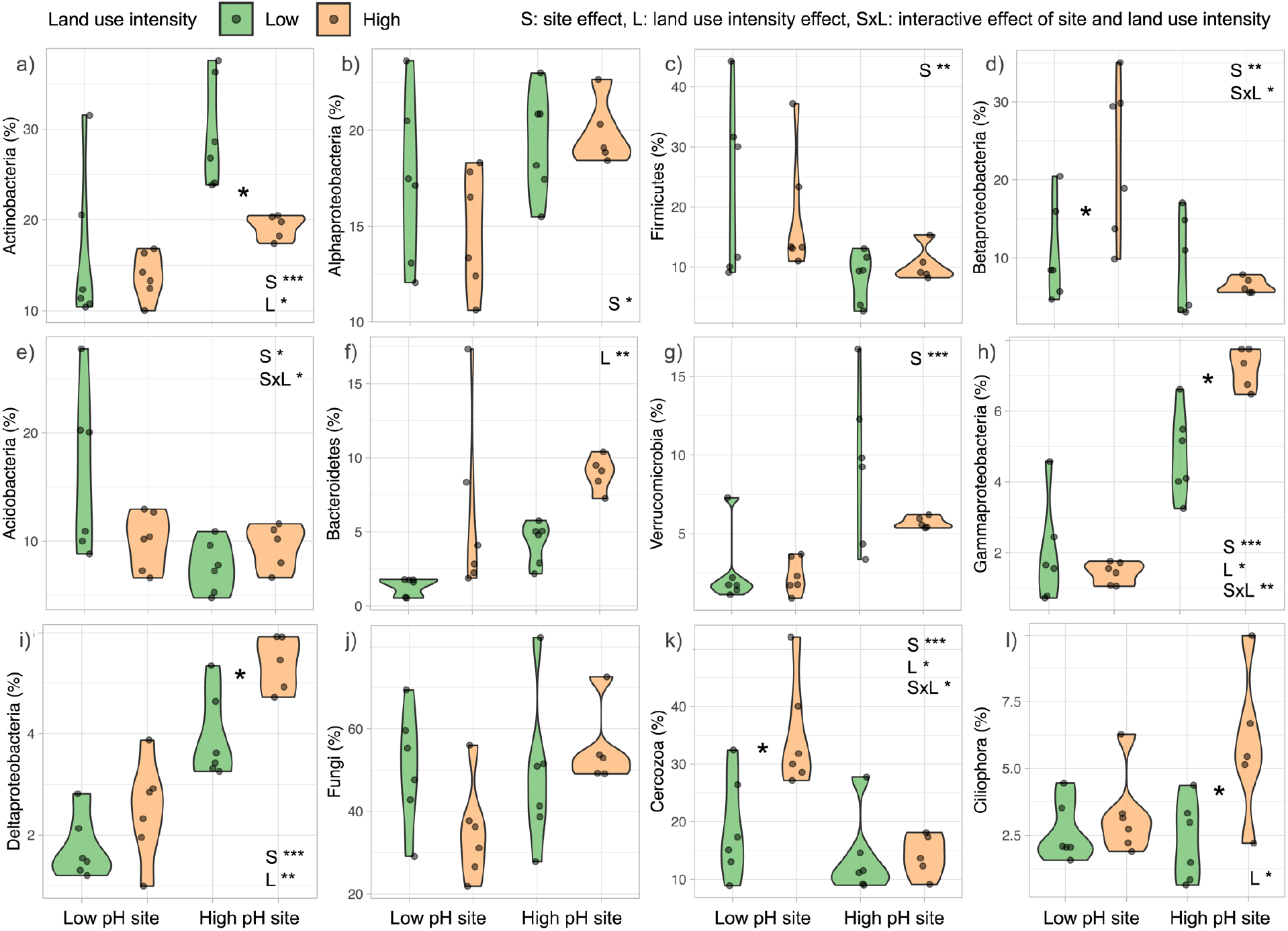
Relative abundance of dominant bacterial and eukaryotic phyla/class: Actinobacteria (a), Alphaproteobacteria (b), Firmicutes (c), Betaproteobacteria (d), Acidobacteria (e), Bacteroidetes (f), Verrucomicrobia (g), Gammaproteobacteria (h), Deltaproteobacteria (i), Fungi (j), Cercozoa (k) and Ciliophora (l). Abundances are displayed across land use intensity treatments: high (orange) and low (green) and the presence of an asterisk between them suggests statistically significant pairwise differences at the site. Also displayed within each plot are statistically significant results of the influencing factors of site (S), land use intensity (L) and their interaction (S×L); *** p < 0.001, ** p < 0.01, * p < 0.05.

The relative abundance of the 12 most abundant taxa (bacteria, fungi and microeukaryotes) were differentially affected by the sites and land use intensity. At the high pH site, low land use intensity soil bacteria were dominated by Actinobacteria, Alphaproteobacteria, and Verrucomicrobia (Fig. 3a,b,g). Land use intensification in high pH soil reduced the abundance of Actinobacteria, but increased that of Gammaproteobacteria, Deltaproteobacteria and Ciliophora (Fig. 3h,i,l). The decline of Actinobacteria under high land use intensity accords with Griffiths et al. [26] who noted that Actinobacteria are common in higher pH soils, but being filamentous, are sensitive to disturbances from agricultural management [58, 59]. Acidobacteria was one of the most dominant bacterial groups in low intensity soils at the low pH site (Fig. 3e). Land use intensification at this site increased the abundance of Betaproteobacteria (from 9% to 26%, Fig. 3d) in accordance with Babin et al. [60], who also observed increases in the abundance of this taxa in improved grassland. Land use intensification at the low pH site also significantly increased the relative abundance of predatory Cercozoa (from 15% to 23%, Fig. 3k).

### Taxa-trait changes due to land use intensification in the high pH site

‘ABC transporters’ were the most abundant protein indicators of low land use at the high pH site. Communities here were associated with a transporter-mediated resource acquisition strategy that is likely more efficient in resource use [25]. The presence of transporters reflects abundant high-quality resource availability, most likely as root exudates and microbial metabolites. The taxonomic assignment of these transporters suggested that they were mostly associated with Alphaproteobacteria (Fig. 4). Although Alphaproteobacteria were not differentially abundant in low land use soils compared to the high land use contrast at this site (Fig. 3b), in terms of the taxonomic distribution of this trait, Alphaproteobacteria were the dominant class differentially expressing this function in low land use soils according to peptide marker abundances relating to this strategy. This implies that members of this class have a resource-uptake optimised strategy in low land use soils that likely contributes to the increase in community-level CUE and therefore promote SOC stabilisation [29].

**Fig. 4:**
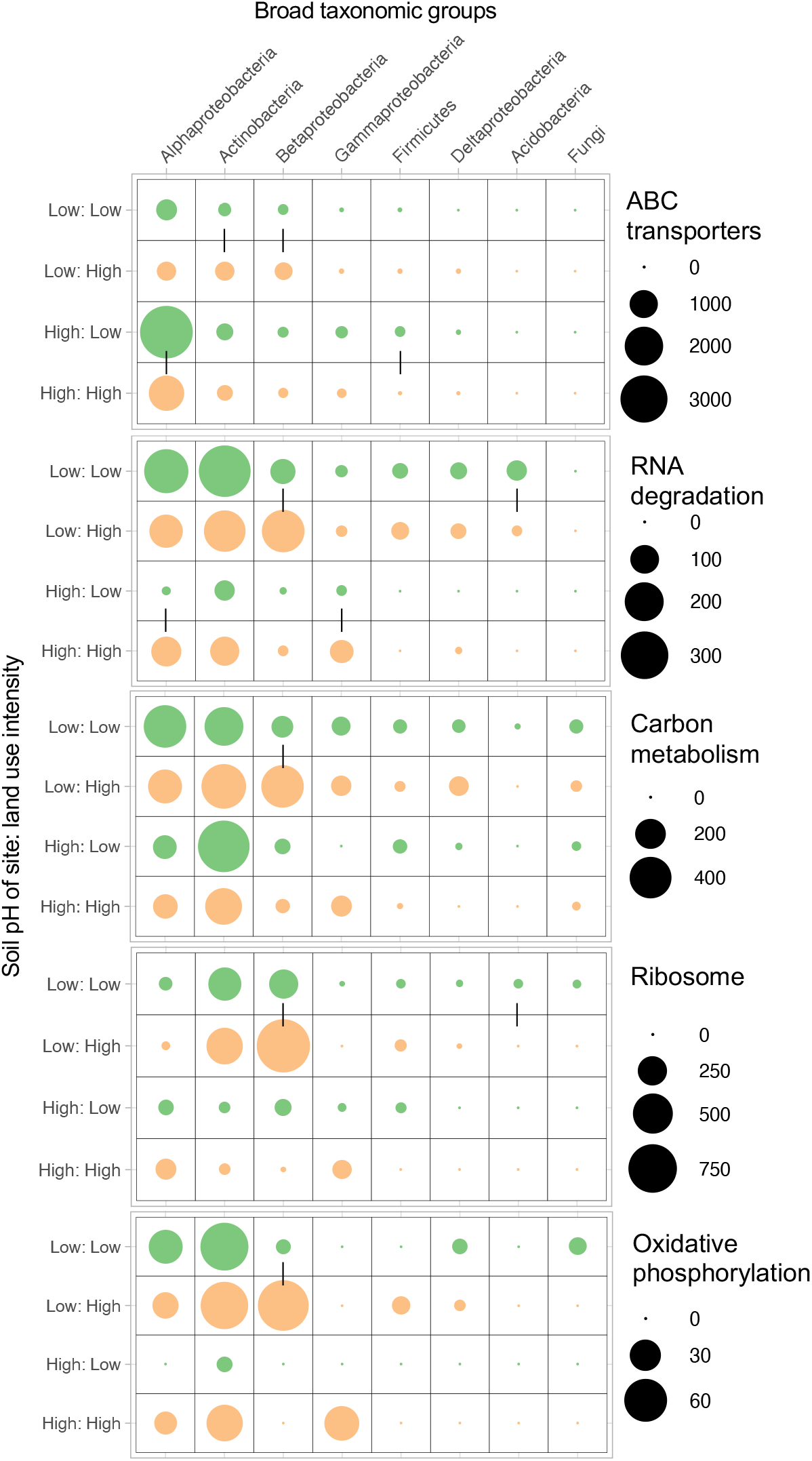
Metaproteomics-derived abundances of functions and their taxonomic lineages were used to link physiological traits to microbial taxa at high (orange) and low (green) land use intensity. Sample labels represent soil pH of the site and land use intensity, for example “Low: Low” means low pH site and low land use intensity. Pairs of circles representing peptide abundances linked by a vertical line within each site are significantly different (p < 0.05).

We observed that land use intensification in high pH soils increased the expression of proteins linked to ‘RNA degradation’ – molecular chaperones that prevent protein aggregation by either re-folding or degrading stress-induced misfolded proteins [29]. Chaperone production in high land use soils indicates microbial investment into stress tolerance. This trait was differentially expressed in the taxa Alphaproteobacteria and Gammaproteobacteria (Fig. 4). Members of these taxa in high land use soils likely excel in a stress tolerance strategy to tide over the dry and disturbed soil conditions.

In addition to the increased expression of stress tolerance traits in Gammaproteobacteria in high land use intensity, proteins linked to ‘oxidative phosphorylation’ (energy generating pathways using ATPase to fuel growth or non-growth maintenance activities) and ‘carbon metabolism’ (central carbon metabolism pathways such glycolysis and TCA cycle) were also differently abundant relative to low land use intensity (Fig. 4). This most likely represents increased energy needs for fast growing taxa with a wasteful metabolism; a life history strategy often associated with copiotrophs such as Gammaproteobacteria that are differentially abundant in high land use soils at this site [25]. We also observed concomitant increased abundance of predatory Ciliophora in high land use soil communities (Fig. 3l), these likely increase in response to the increased abundance of their bacterial prey – a hypothesis that needs testing. These microbivorous protists could contribute to SOC stabilisation directly through increased necromass contributions, but also through their influence on the assemblage and function of the microbiome [61].

### Taxa-trait changes due to land use intensification in the low pH site

Stress proteins were the most abundant protein indicators at the low pH site in both low and high land use soils but with higher relative abundances in the low intensity land use treatment [25]. This is likely a physiological response to the acidic and wet conditions in the low intensity soils through increased expression of ‘RNA degradation’ proteins such as Chaperonin GroEL and molecular chaperone DnaK. They were differentially abundant in the phylum Acidobacteria in the low intensity relative to high intensity soils (a dominant taxonomic group in low intensity soils at the low pH site; Fig. 3a). But there were other taxa that also had higher expression of this trait in the low intensity soils.

Land use intensification at the low pH site increased the abundance of Betaproteobacteria (from 9% to 26%, Fig. 3d) in accordance with Babin et al. [60], who also observed increases in the abundance of this taxa in improved grassland. Members of the class Betaproteobacteria also showed differentially abundant stress proteins in high intensity soils. However, they also showed increased abundance of ‘ABC transporters’ (Fig. 4), indicating an uptake-optimised resource acquisition strategy reflecting the increased abundance of resources under high intensity land use. This is a result of alleviation of constraints on microbial decomposition of organic matter due to increase in pH and decrease in wetness and anoxia. Betaproteobacteria also had increased expression of proteins linked to ‘carbon metabolism’, ‘ribosome’, and ‘oxidative phosphorylation’ pathways; a land use intensification response very similar to that of Gammaproteobacteria in high pH soils. This response likely represents a shift towards increased growth and turnover in a stressed and disturbed environment. Land use intensification also significantly increased the relative abundance of predatory Cercozoa (from 15% to 23%, Fig. 3k) at low pH, that may be responding to increased prey availability under high intensity land use, such as the increase in fast-growing Betaproteobacteria. The increase of Cercozoa under high land use intensity at the low pH site mirrors the increase of Ciliophora under high land use intensity at the high pH site, which suggests that the dominance of distinct bacterial groups might be associated with distinct predatory protozoan groups driving turnover of carbon to a variable degree.

### Taxa-trait changes related to mechanisms of soil carbon cycling

The observed shifts in trait-taxa linkages are in line with our hypothesis that land use intensification leads to shifts in microbiome structure and its associated traits that has consequences for ecosystem CUE. In our high pH site, low intensity land use with no resource limitation and minimum stress resulted in a microbial community that is dominated by taxonomic sub-groups within Alphaproteobacteria that have an efficient transporter-mediated resource-uptake optimised life history strategy with limited investment in stress tolerance traits. This likely increased the microbial biomass (and therefore necromass production) promoting SOC stabilisation pathways. However, increased land use intensification in high pH soils caused resource limitation and stress in microbes. This led to proliferation of microbial sub-groups within Alphaproteobacteria and Gammaproteobacteria that likely excel in an inefficient stress-tolerance life history strategy diverting resources away from biosynthesis and necromass formation and resulted in increased carbon loss and reduced SOC stabilisation. The taxa-trait linkages were vastly different in low pH soils. Here, soils under low intensity land use were dominated by Acidobacteria excelling in stress tolerance traits highlighting a life history strategy that is adapted to the acidic, wet, and anoxic soil conditions. The low growth rates observed in these soils suggest lower rates of decomposition and accumulation of undecomposed plant organic matter. Increased land use intensification in these low pH soils reduced soil acidity, wetness and anoxia which led to increased microbial growth likely due to alleviation of microbial physiological constraints. This results in a shift towards Betaproteobacteria excelling in stress tolerance and resource acquisition strategies that fuel their higher growth rates which could be linked to increased decomposition and loss of the historically accumulated SOC.

Our research reveals that land use intensification induced shifts in microbial taxa and their life history strategies that were pH dependant. Changes in soil characteristics selects for a new community with different traits (environmental filtering) rather than the community shifting its physiology (phenotypic adaptation) [4, 32]. Our study accords with previous trait-based approaches that have demonstrated that microbial efficiency declines along gradients of environmental stress, as increased stress through altitude [34] and salinity [36] results in increased stress tolerance and resource acquisition life history strategies that reduce microbial CUE and negatively influence the microbially-derived SOC formation. Further, our findings of increased abundance of predatory protozoa in response to increased land use intensification, could be crucial for carbon turnover and food web connectivity. This is especially pertinent, as protists are known to be key for promoting the formation of necromass and consequently more persistent mineral-associated organic matter [58].

Here we successfully used a trait-based framework to link taxonomic information to traits and rates of carbon cycling in soils. In this sense, this approach encompasses many of the concepts required to envisage soil health [62], by focussing on the function of the active microbiome and its emergent traits (such as CUE) but also on other biogeochemical factors that are key to determining the balance of SOC decomposition and stabilisation pathways. We also demonstrate how CUE-SOC relationship can be decoupled and how variable pathways of decomposition and stabilisation of particulate and mineral associated organic matter can influence SOC loss or gain in response to land use change. This holistic understanding will be fundamental to predict soil’s ability to recover from the combined stressors of intensification along with environmental change, to ensure that our soils and their microbiomes remain resilient and productive under global change [63].

## Acknowledgements

This work was funded by a UK Natural Environment Research Council NERC sponsored Daphne Jackson Trust Fellowship awarded to LC, the European Union’s Horizon 2020 research and innovation program under the Marie Skłodowska–Curie grant no. 655240 awarded to AAM and the UK Natural Environment Research Council under a Soil Security Programme grant (NE/M017125/1) to RIG. We also wish to thank Kate Buckeridge and Kelly Mason for soil sampling, and Lara Oudot and Emily MacDonald for technical support.

## Author contributions

AAM conceived, designed, and carried out the experiment and laboratory analyses; TG performed amplicon sequencing; NJ performed metaproteomic analysis; GG and RIG contributed new reagents and analytical tools; AAM and LC performed statistical analyses; LC drafted the manuscript with supervision from AAM and all authors were involved in critical revision and approval of the final version.

